# Immune Landscape of Invasive Ductal Carcinoma Tumour Microenvironment Identifies a Prognostic and Immunotherapeutically Relevant Gene Signature

**DOI:** 10.1101/620849

**Authors:** Xuanwen Bao, Run Shi, Kai Zhang, Shan Xin, Xin Li, Yanbo Zhao, Yanfang Wang

## Abstract

**Background:** Invasive ductal carcinoma (IDC) is a clinically and molecularly distinct disease. Tumour microenvironment (TME) immune phenotypes play crucial roles in predicting clinical outcomes and therapeutic efficacy.

**Method:** In this study, we depict the immune landscape of IDC by using transcriptome profiling and clinical characteristics retrieved from The Cancer Genome Atlas (TCGA) data portal. Immune cell infiltration was evaluated via single-sample gene set enrichment (ssGSEA) analysis and systematically correlated with genomic characteristics and clinicopathological features of IDC patients. Furthermore, an immune signature was constructed using the least absolute shrinkage and selection operator (LASSO) Cox regression algorithm. A random forest algorithm was applied to identify the most important somatic gene mutations associated with the constructed immune signature. A nomogram that integrated clinicopathological features with the immune signature to predict survival probability was constructed by multivariate Cox regression.

**Results:** The IDC were clustered into low immune infiltration, intermediate immune infiltration, and high immune infiltration by the immune landscape. The high infiltration group had a favourable survival probability compared with that of the low infiltration group. The low-risk score subtype identified by the immune signature was characterized by T cell-mediated immune activation. Additionally, activation of the interferon-α response, interferon-γ response and TNF-α signalling via the NFκB pathway was observed in the low-risk score subtype, which indicated T cell activation and may be responsible for significantly favourable outcomes in IDC patients. A random forest algorithm identified the most important somatic gene mutations associated with the constructed immune signature. Furthermore, a nomogram that integrated clinicopathological features with the immune signature to predict survival probability was constructed, revealing that the immune signature was an independent prognostic biomarker. Finally, the relationship of VEGFA, PD1, PDL-1 and CTLA-4 expression with the immune infiltration landscape and the immune signature was analysed to interpret the responses of IDC patients to immune checkpoint inhibitor therapy.

**Conclusion:** Taken together, we performed a comprehensive evaluation of the immune landscape of IDC and constructed an immune signature related to the immune landscape. This analysis of TME immune infiltration patterns has shed light on how IDC respond to immune checkpoint therapy and may guide the development of novel drug combination strategies.

## Introduction

Invasive ductal carcinoma (IDC) is a clinically and molecularly distinct disease. IDCs are typically of high histologic grade and high mitotic index. HER2 overexpression or amplification is detected in 20% of these tumours (1). IDC tends to metastasize to bone, liver, and lung, whereas invasive lobular carcinoma (ILC) has a higher tendency to metastasize in gastrointestinal and genital tracts, serosal cavities, and meninges (2). IDCs usually form glandular structures in contrast to the small clusters formed by ILCs. The loss of CDH1 leads to the discohesive morphology in ILCs, whereas IDCs maintain intact cell adhesion (3). Furthermore, the frequency of recurrently mutated genes and recurrent copy-number alterations often differs significantly between IDCs and ILCs (3). These features are generally associated with a poor prognosis. Taken together, these differences suggest that ILCs and IDCs are distinct cancer types and progress along different pathways.

Genetic and epigenetic changes contribute to the progression of tumour progression and recurrence in different cancer types. However, accumulated evidence indicates that the tumour microenvironment (TME) has clinicopathological significance in predicting survival outcomes and assessing therapeutic efficacy factors (4, 5). TME cells constitute a vital element of cellular and noncellular components in the tumoural niche, including extracellular matrix and cellular components, such as fibroblasts, adipose cells, immune-inflammatory cells, and neuroendocrine cells. Previous studies have revealed that immune cells in the TME modulate cancer progression and are attractive therapeutic targets (6, 7). To date, the comprehensive landscape of immune cells infiltrating the TME of IDCs has not yet been elucidated. We propose that IDCs have a distinct immune landscape and that the immune landscape might lead to different prognoses and treatment responses. In this study, by applying several computational algorithms, we estimated the abundance of immune cells in the TME of IDCs and analysed the correlation of the immune landscape with genomic characteristics and pathological features of IDCs. Furthermore, we built an immune signature based on the TME immune phenotype, which is a robust prognostic biomarker and predictive factor for the response to immune-checkpoint inhibitors.

## Method

### Data download

TCGA RNA-seq datasets and clinical data for IDCs were downloaded by UCSC Xena browser (https://xenabrowser.net/). GSE20685 and GSE86948 were downloaded from the Gene Expression Omnibus (GEO) database.

### Implementation of Single-Sample Gene Set Enrichment Analysis (ssGSEA)

We obtained the marker gene set for immune cell types from Bindea et al (8). We used the ssGSEA program to derive the enrichment scores of each immune-related term. In brief, the infiltration levels of immune cell types were quantified by ssGSEA in the R package gsva (9). The ssGSEA applies gene signatures expressed by immune cell populations to individual cancer samples. The computational approach used in our study included 24 immune cells types that are involved in innate immunity and adaptive immunity. Tumours with qualitatively different immune cell infiltration patterns were grouped using hierarchical agglomerative clustering (based on Euclidean distance and Ward’s linkage).

The T cell infiltration score (TIS) was defined as the average of the standardized values for CD8+ T, central memory CD4+ T, effector memory CD4+ T, central memory CD8+ T, effector memory CD8+ T, Th1, Th2, Th17, and Treg cells. The obtained cytotoxic activity scores (CYT) score was calculated by the geometrical mean of PRF1 and GZMA (10). The CD8+ T/Treg ratio was the digital ratio of the ssGSEA scores for these two cell types. The correlation between risk score and immune cell ssGSEA score was calculated by Pearson correlation.

### LASSO regularization

LASSO (least absolute shrinkage and selection operator) is an important regularization in many regression analysis methods (e.g., COX regression and logistic regression) (10). The idea behind LASSO is that an L1-norm is used to penalize the weight of the model parameters. Assuming a model has a set of parameters, the LASSO regularization can be defined as:

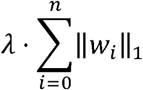

It can also be expressed as a constraint to the targeted objective function:

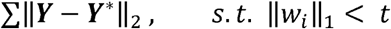

An important property of the LASSO regularization term is that it can force the parameter values to be 0, thus generating a sparse parameter space, which is a desirable characteristic for feature selection. In our analysis, 19 genes which highly associated with OS were used as the input.

### Differentially expressed gene (DEG) analysis

DEG analysis was performed by the Limma package (11). The samples were separated into a high-risk score group and a low-risk score group. An empirical Bayesian approach was applied to estimate the gene expression changes using moderated t-tests. The Q value (adjusted p value) for multiple testing was calculated using the Benjamini-Hochberg correction. The DEGs were defined as genes with a Q value less than 0.05. The clusterProfiler R package was applied for the GO analysis (12). GSEA was applied with the GSEA software.

### Co-expression gene network based on RNA-seq data

The Weighted correlation network analysis (WGCNA) was used to construct the gene co-expression network (13). The co-expression similarity *s*_*i,j*_ was defined as the absolute value of the correlation coefficient between the profiles of nodes *i* and *j*:

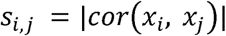

where *x*_*i*_ and *x*_*j*_ are expression values of for genes *i* and *j*, and *s*_*i,j*_ represent Pearson’s correlation coefficients of genes *i* and *j*, respectively.

A weighed network adjacency was defined by raising the co-expression similarity to a power *β*:

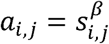

with *β*≥1. We selected the power of *β* = 5 and scale-free R^2^ = 0.95 as the soft-thresholding parameters to ensure a signed scale-free co-expression gene network. Briefly, network construction, module detection, feature selection, calculations of topological properties, data simulation, and visualization were performed. Modules were identified via hierarchical clustering of the weighting coefficient matrix. The module membership of node *i* in module *q* was defined as:

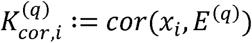

where *x*_*i*_ is the profile of node *i*, and E(*q*) is the module eigengene (the first principal component of a given module) of module *q*. The module membership measure 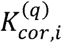, lies in [−1,1] and specifies how close node *i* is to module *q, q* = 1,…,.*Q*.

By evaluating the correlations between the immune infiltration status, immune signature of IDCs and the module membership of each module, a brown module was selected for further analysis.

### Statistical analysis

A random forest algorithm was applied to find the most important somatic mutation associated with the immune signature. Survival outcome analysis modelled the results in reference to the patient OS and RFS. P-values and Hazard ratios were obtained from univariate Cox proportional-hazards regression models using the R package survival. Multivariate Cox regression was used to calculate the coefficients in the nomogram. The nomogram was plotted by the rms package. The time-dependent AUC value was calculated by the survivalROC package.

## Results

### Immune Phenotype Landscape in the TME of IDC

Immune cell populations modulate diverse immune responses and lead to antitumour effects by infiltrating the IDC TME. The immune cell infiltration status was assessed by applying the ssGSEA approach to the transcriptomes of IDCs. Twenty-four immune-related terms were incorporated to deconvolve the abundance of diverse immune cell types in IDCs. The IDCs were clustered into 3 clusters (low infiltration: 208; intermediate infiltration: 430; and high infiltration: 130) in terms of immune infiltration by applying an unsupervised clustering algorithm (Fig. 1A). By applying hierarchical cluster analysis and K-means clustering analysis, we constructed a TME cell network, depicting a comprehensive landscape of tumour-immune cell interactions and their effects on the OS of patients with IDC (Figs. 1B, S1 and S2). The TME immune cells were clustered into 4 clusters, and the correlation among different immune cell types is shown in Fig. 1B. The association of OS and RFS with different clusters of IDCs was analysed by a pairwise log-rank test. The results indicated that the high infiltration group had a favourable survival probability compared with that of the low infiltration group (Fig. 1C and 1D).

**Fig 1.**
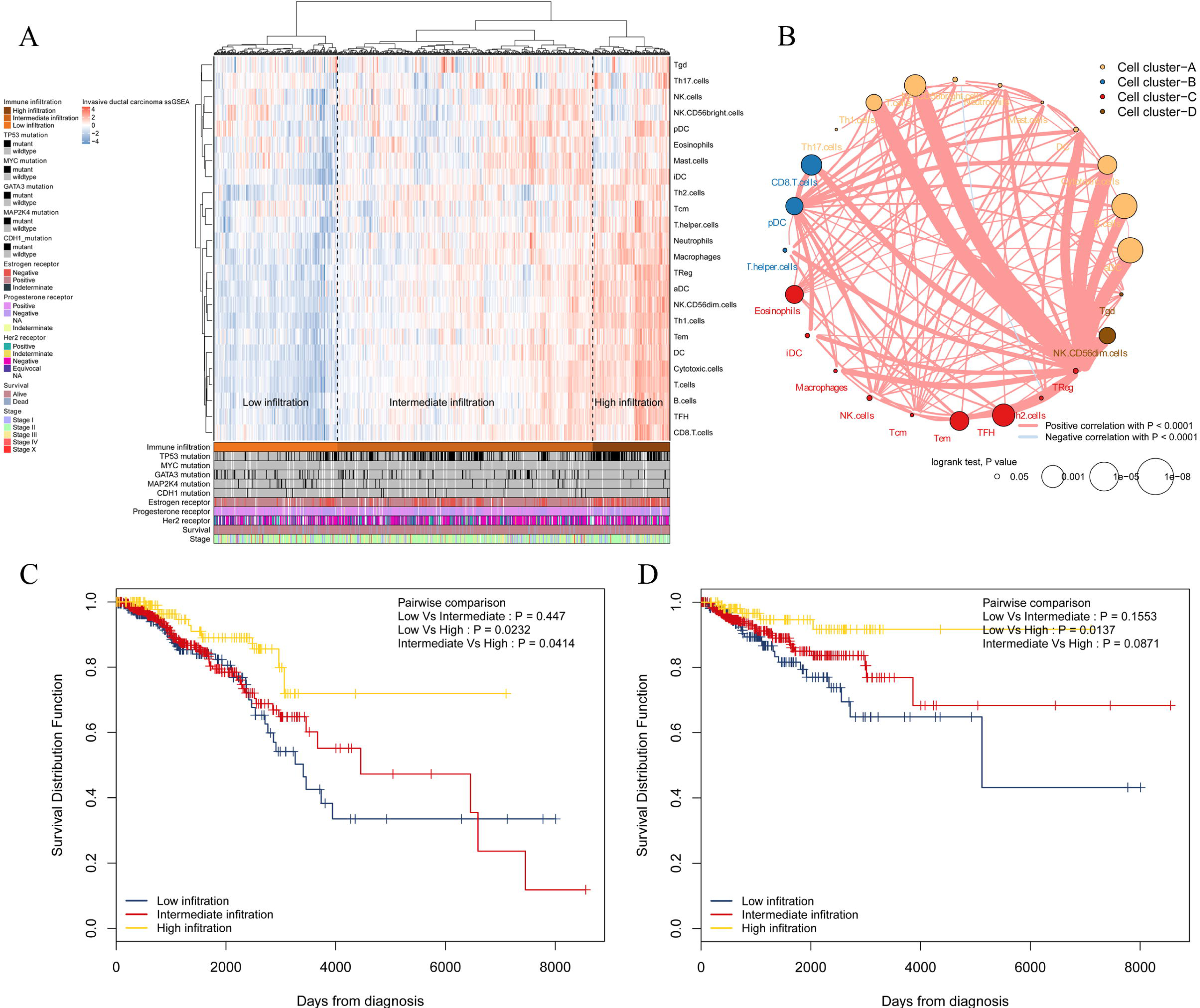
Immune landscape of IDCs and the TME characteristics. A, Unsupervised clustering of IDC patients from the TCGA cohort using ssGSEA scores from immune cell types. Mutation status of TP53, MYC, GATA3, MAP2K4, and CDH1, status of the oestrogen receptor, status of the progesterone receptor, status of Her2, survival, and stage are shown as patient annotations in the lower panel. Hierarchical clustering was performed with Euclidean distance and Ward linkage. Three distinct immune infiltration clusters, here termed high infiltration, median infiltration, and low infiltration, were defined. B, Interaction of the TME immune cell types. The size of each term represents the survival impact of each TME cell type, calculated by log_10_ (log-rank test P value). The connection of TME immune cells represents interactions between both. The thickness of the line indicates the strength of the correlation calculated by Spearman correlation analysis. Positive correlations are represented in red, and negative correlations are represented in blue. The immune cell cluster was clustered by the hclust method. Immune cell cluster-A, yellow; cell cluster-B, blue; cell cluster-C, red; and cell cluster-D, brown. C, Kaplan-Meier curves for OS of IDC patients showing that the high immune infiltration group had a favourable outcome compared with the other groups. D, Kaplan-Meier curves for RFS of IDC patients showing that the high immune infiltration group had a favourable outcome compared with other groups. IDC: invasive ductal carcinoma; TME: tumour microenvironment; TCGA: The Cancer Genome Atlas; OS: overall survival; and RFS: recurrence-free survival.

### Construction of the immune signature

A total of 413 genes were involved in the 24 immune-related terms. We applied the univariate COX regression based on the survival datasets of patients with IDC and the expression profiles of the 413 genes. The 19 most significant genes were selected with the criteria of a p value less than 0.0005 (Fig. 2A). The expression profiles of the 19 genes are shown in Fig. 2B. LASSO Cox regression was performed on the 19 genes to identify the most important features in terms of predicting the survival of IDC patients (Fig. 2C, 2D and 2E). By forcing the sum of the absolute value of the regression coefficients to be less than a fixed value, certain coefficients were reduced to exactly zero, and the most powerful prognostic features (QRSL1, TIMM8A, IGHA1, BATF, KLRB1, SPIB, and FLT3LG) were identified with relative regression coefficients. Cross-validation was applied to prevent over-fitting. A 7-gene immune signature was constructed according to the individual coefficients of the genes. Then, we calculated the risk score for each IDC patient and ranked them (Fig. 2F). Fig. 2G shows the survival overview in the IDC patients. A heatmap showed that patients in the high-risk group tended to have increased QRSL1 and TIMM8A expression levels, as well as decreased expression levels of IGHA1, BATF, KLRB1, SPIB, and FLT3LG (Fig. 2G). The Kaplan-Meier curve and Cox regression suggested that patients with high risk scores had significantly worse OS and RFS than those with low risk scores (HR=2.94, p<0.0001 and HR=2.28, p=0.001, respectively) (Fig. 2H and 2I). The effect of the 7 genes on the OS and RFS of IDC patients is shown in Fig. S3 and Fig. S4, respectively. To confirm our findings in the IDC cohort, we validated the prognostic function of the immune signature in two independent GEO cohorts (GSE20685 and GSE86948). The risk score was calculated for each patient by using the same formula as in the IDC cohort. The GSE20685 and GSE86948 cohorts were used to predict the OS of BRCA patients based on our immune signature model. Consistent with our previous findings, the Kaplan-Meier curve suggested a significantly better overall survival in the low-risk group than in the high-risk group (Fig. S5A and S5B).

**Fig 2.**
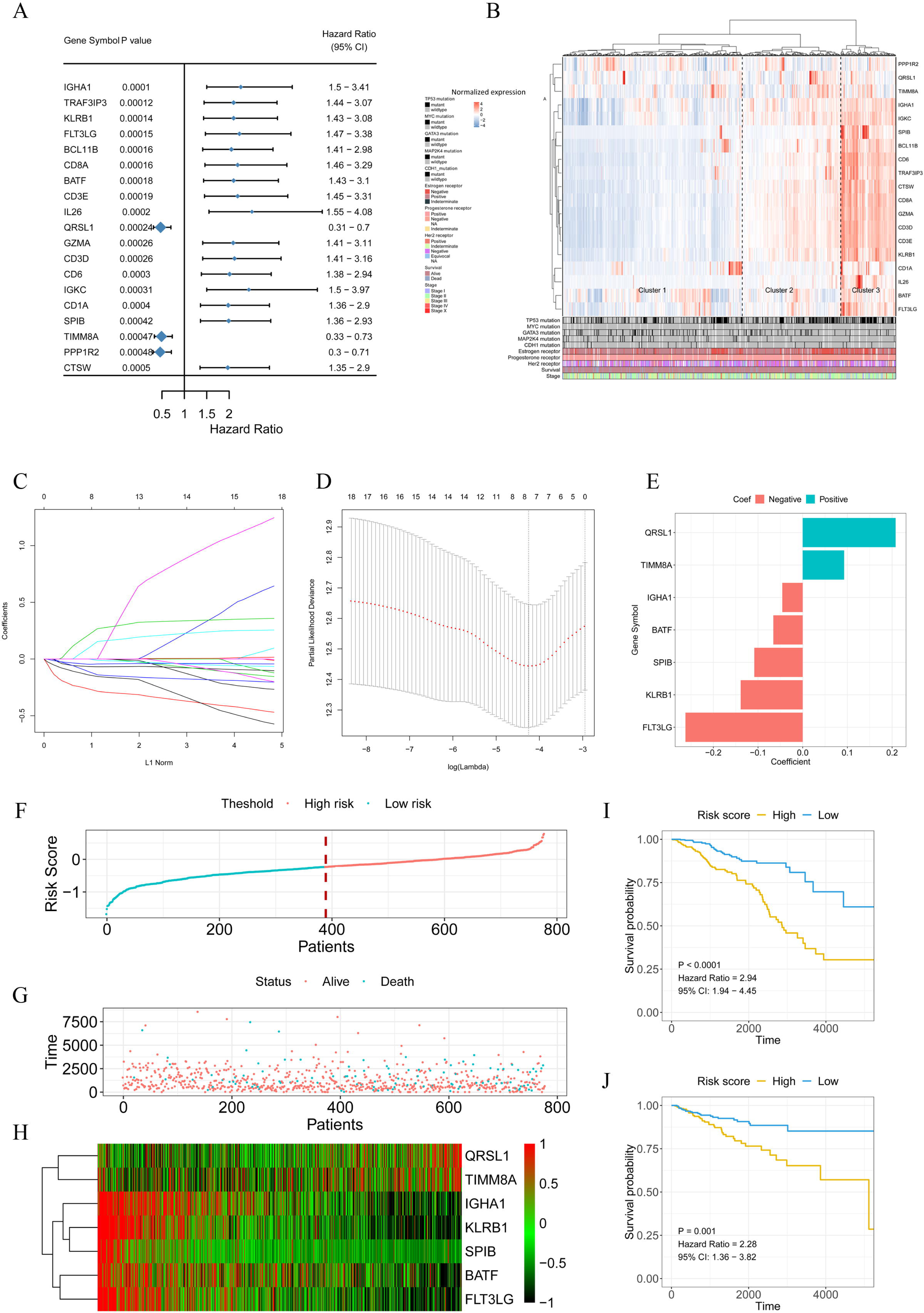
Signature-based risk score is a promising marker of survival in IDC patients. A, The HR and P value from the univariable Cox HR regression of selected genes in the immune terms (Criteria: P value < 0.001). B, The expression of the selected genes shown by heatmap. Mutation status of TP53, MYC, GATA3, MAP2K4, and CDH1, status of the oestrogen receptor, status of the progesterone receptor, status of Her2, survival, and stage are shown as patient annotations in the lower panel. Hierarchical clustering was performed with Euclidean distance and Ward linkage. C and D, LASSO Cox analysis identified 7 genes most correlated with overall survival, and 10-round cross validation was performed to prevent overfitting. E, Coefficient distribution of the gene signature. F, Risk score distribution. G, Survival overview. H, Heatmap showing the expression profiles of the signature in the low- and high-risk groups. I, Patients in the high-risk group exhibited worse OS than those in the low-risk group. J, Patients in the high-risk group exhibited worse RFS than those in the low-risk group. IDC: invasive ductal carcinoma; OS: overall survival; and RFS: recurrence-free survival.

### The low risk score was associated with active infiltration status and high cytotoxic potential

High infiltration status showed a lower risk score than the intermediate infiltration status and low infiltration status showed (Fig. 3A). Similarly, patients with a low risk score had a higher proportion of high immune infiltration than patients with a high risk score (Fig. 3B). The presence of high immune infiltration in patients was linked to a low risk score and was associated with a favourable outcome (Fig. 3C). To compare cytotoxic function with the immune landscape and immune signature that we constructed, the associated signatures were identified for each patient. IDCs with high infiltration status and low risk score were associated with increased levels of immune activation. The TIS (p < 0.0001 and p < 0.0001, respectively) (Fig. 3D and 3H), interferon-γ signature (p < 0.0001 and p < 0.0001, respectively) (Fig. 3E and 3I), and CYT (p < 0.0001 and p < 0.0001, respectively) (Fig. 3F and 3J) were increased in the low-risk score group and high infiltration group. The ssGSEA score of DCs was higher in the low-risk score group than in the high-risk score group. The Kaplan-Meier curve showed that in the low-risk score group, the ssGSEA score of DC cells affected survival but did not affect the high-risk score group (Fig. S6A, S6B and S6C). These data indicate that compared with high-risk score tumours, low-risk score tumours have a distinct immune phenotype, characterized by increased immune infiltration and increased levels of immune activation.

**Fig 3.**
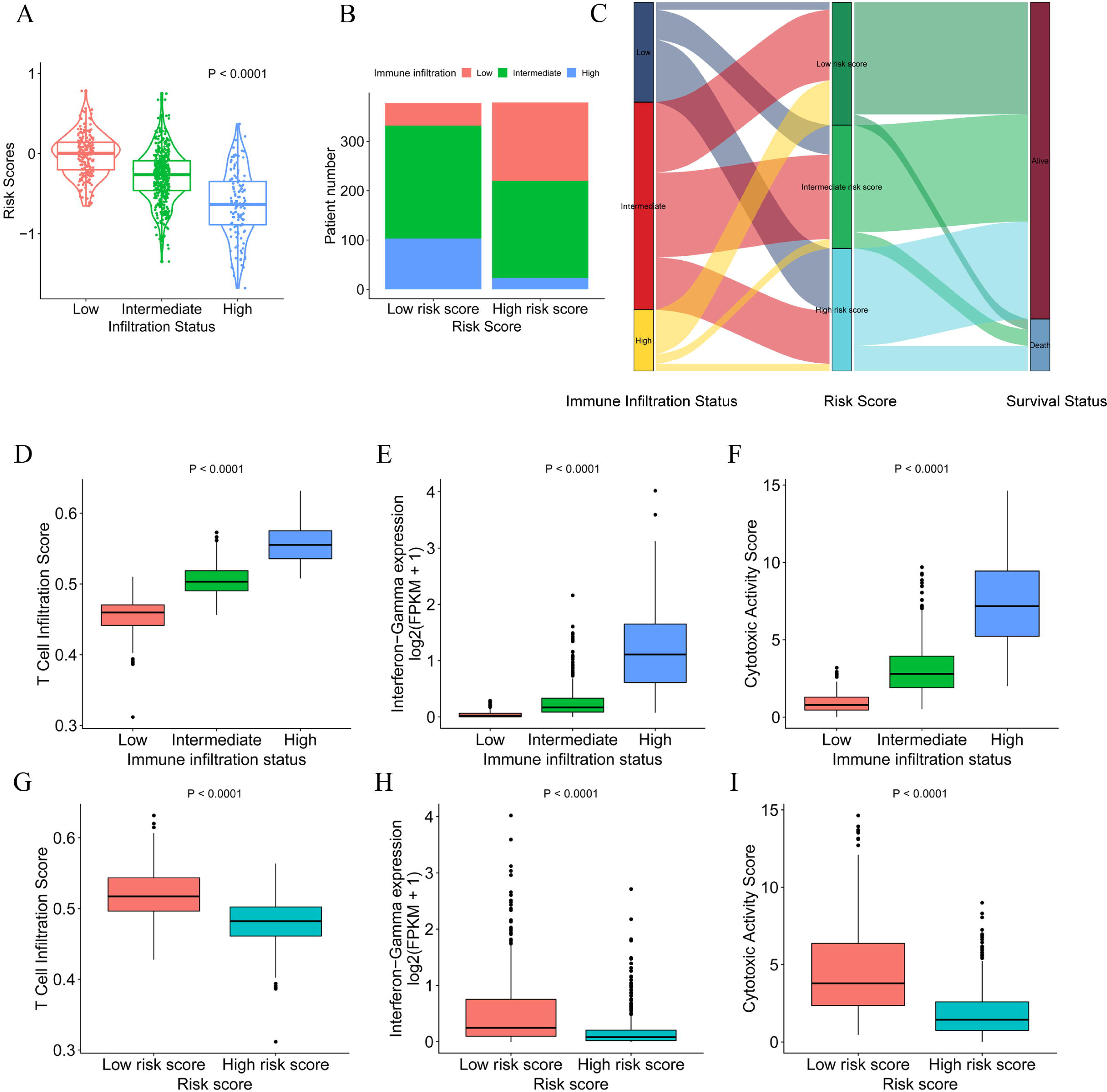
Heterogeneous immune cell infiltration in the low- and high-risk score groups. A, The distribution of risk scores in low, mediate and high immune infiltration patterns. B, The distribution of immune infiltration patterns in the low- and high-risk score groups. C, Alluvial diagram of immune infiltration patterns in groups with different risk scores and survival outcomes. D, TIS in low, mediate and high immune infiltration patterns. E, Relative interferon-γ signature in low, mediate and high immune infiltration patterns. F, Comparison of relative CYT in low, mediate and high immune infiltration patterns. G, Relative TIS in the low- and high-risk score groups. E, Relative interferon-γ signature in the low- and high-risk score groups. F, Comparison of relative CYT in the low- and high-risk score groups. TIS: T cell infiltration score; CYT: cytotoxic activity scores.

### The low-risk score was associated with increased T cell infiltration

The association of risk score and immune-related cells was analysed by Pearson correlation. Cytotoxic cells, CD8+ T cells, T cells and the 6 other most significant immune-related cell types are shown in Fig. 4. A high level of correlation was found between the risk score and the T cell-mediated immune response. The ssGSEA scores of 24 immune-related terms in the low, intermediate, and high immune status and low- and high-risk score groups are shown in Fig. S7A and S7C. The p value and difference in the mean ssGSEA score from the high- and low-infiltration status and low- and high-risk score groups are shown in Fig. S7B and Fig. S7D. The proportions of low, intermediate, and high immune infiltration status in different pathological subtypes and different AJCC stages of IDC are shown in Fig. S7E and Fig. S7F. The triple-negative subtype of IDCs had a higher proportion of high infiltration status IDCs than other pathological subtypes, indicating an active immune response in triple-negative IDCs. The risk score distribution in different pathological subtypes and different AJCC stages of IDC are shown in Fig. S5G and Fig. S5H. The luminal A subtype had a lower risk score than the other pathological subtypes.

**Fig 4.**
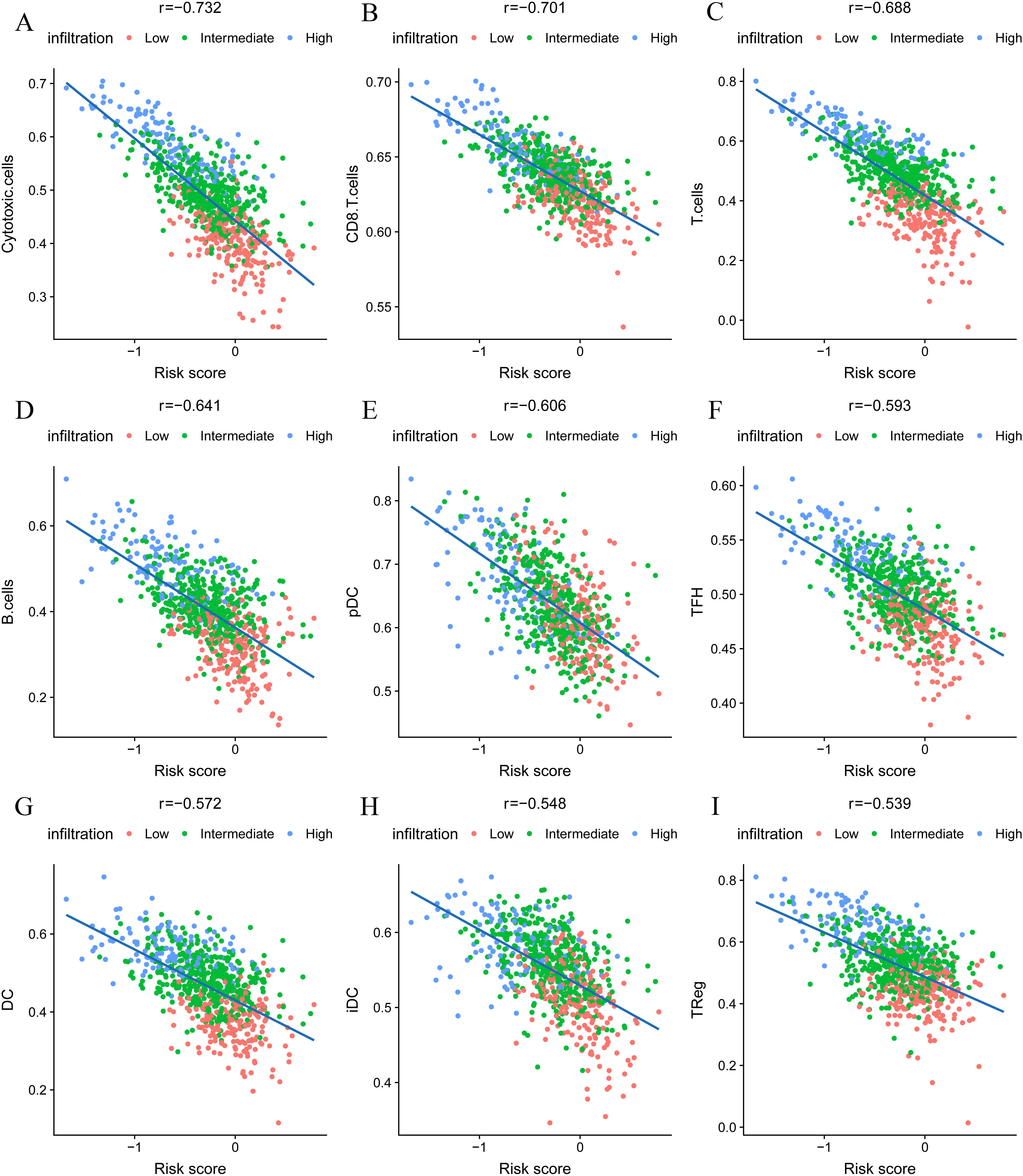
The nine most significant correlations of risk score with immune cell infiltration ssGSEA score.

### Functional annotation and WGNCA of the transcriptomes of IDC patients

To identify the underlying biological characteristics of the constructed immune signature, DEG analysis was performed based on the high-risk score group and low-risk score group. The heatmap depicts the significant DEGs between the two groups (Fig. 5A). The GO analysis indicated that T cell activation, positive regulation of leukocyte cell-cell adhesion, and regulation of lymphocyte activation were the most significantly enriched biological processes between the high-risk score group and the low-risk score group (Fig. 5B). The GSEA results showed that allograft rejection, IL-6/JAK/STAT3 signalling, the inflammatory response, interferon-α response, interferon-γ response and TNF-α signalling via the NFκB pathway were the predominant upregulated pathways in the low-risk score group. In contrast, the E2F targets, G2M checkpoints, MTORC1 signalling and protein secretion pathways were significantly downregulated in the low-risk score group (Fig. 5C and 5D). To further identify the underlying biological characteristics in the immune signature, WGCNA was performed, and the correlation of risk score and immune infiltration status with module membership was analysed. The soft threshold selection is shown in Fig. S8. The module-trait heatmap illustrates that the brown module had a significant p value with both immune signature and immune infiltration status (Fig. 5E); the coefficients were −0.64 and 0.8, respectively. The association between module membership and gene significance for each gene in the brown module is shown in Fig. 5F. The genes from the brown module with a coefficient greater than 0.5 were selected as hub genes, and GO enrichment analysis revealed that T cell activation and lymphocyte activation were the most significantly enriched biological processes, which further confirmed the results from the DEG analysis.

**Fig 5.**
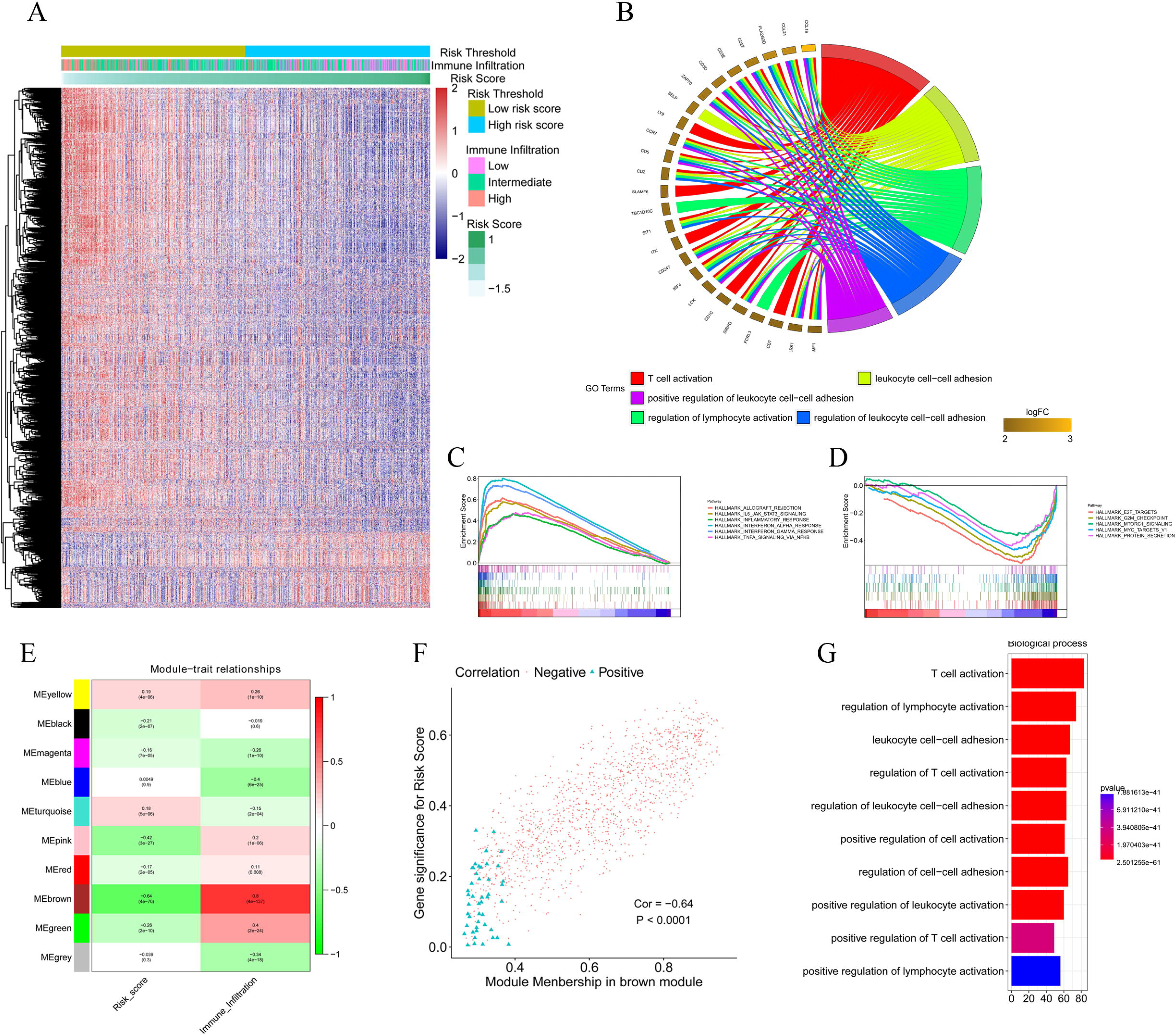
Functional annotation of the immune signature and WGCNA of the IDC transcriptome. A, Heatmap showing the transcriptome expression profiles of the low- and high-risk groups. B, GO analysis based on the significant genes in the comparison between low- and high-risk groups. C and D, GSEA revealed that most significant hallmarks correlated with the immune signature. E, Correlation between modules and traits. G, The correlation between module membership and gene significance in the brown module. H, GO analysis based on the hub genes in the brown module. GO: gene ontology; GSEA: gene set enrichment analysis.

### Mutation load and immune signature

The spectrum of somatic mutations in patients with IDCs is known to be varied. We next investigated the distributions of somatic mutations and observed different patterns among IDCs in terms of total mutation burden (TMB). The risk score from the immune signature had a positive correlation with TMB in IDC patients (Fig. 6A). By applying a random forest algorithm, we identified 35 highly variable mutated genes that were associated with the immune signature (Fig. 6B). TP53 was the predominant gene of the 35 identified genes.

**Fig 6.**
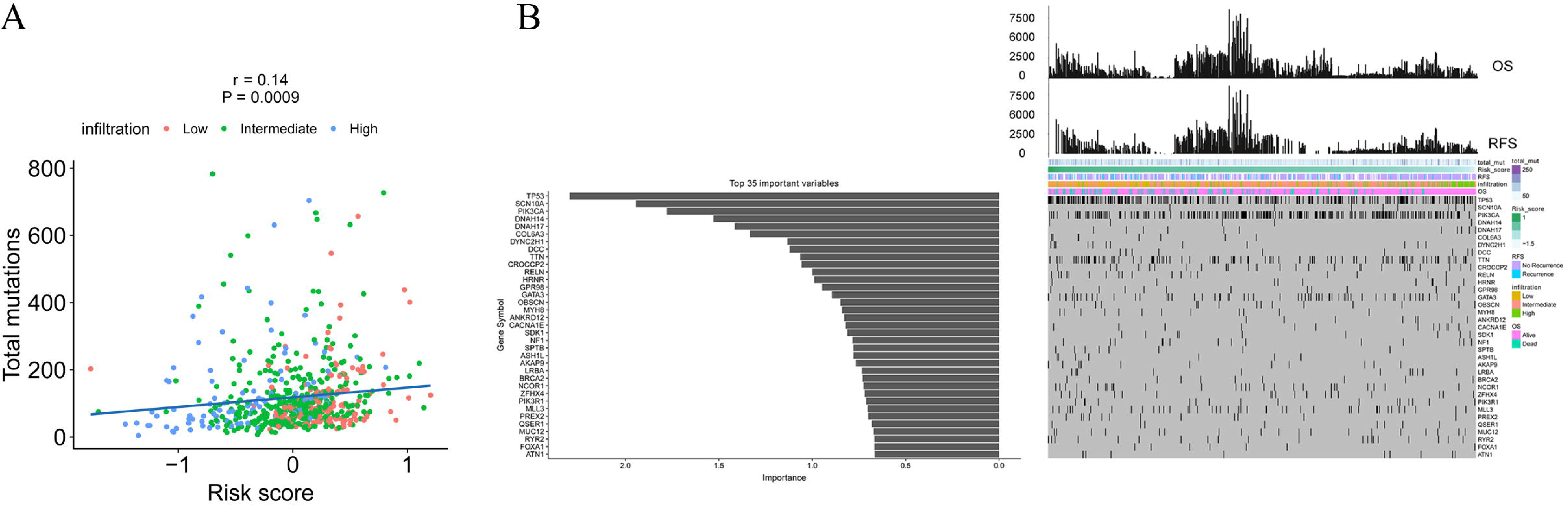
The association of the immune signature with cancer somatic mutations. A, The correlation between the immune signature and IDC somatic mutations. B, Distribution of somatic mutations correlated with the immune signature. The upper bar plot indicates OS and RFS per patient, whereas the left bar plot shows the importance of the somatic mutations correlated with the immune signature. IDC: invasive ductal carcinoma; OS: overall survival; and RFS: recurrence-free survival.

### Construction of a nomogram to predict overall survival in IDC patients

We constructed a nomogram that integrated clinicopathological features with the immune signature to predict the survival probability of IDC patients (Fig. 7A). The AUC(t) functions of the multivariable models were developed to indicate how well these features serve as prognostic markers. Compared to other features, such as signature-based risk score, AJCC-TNM stage and total mutation burden, the nomogram consistently showed the highest predictive power for overall survival in the follow-up period (Fig. 7B).

**Fig 7.**
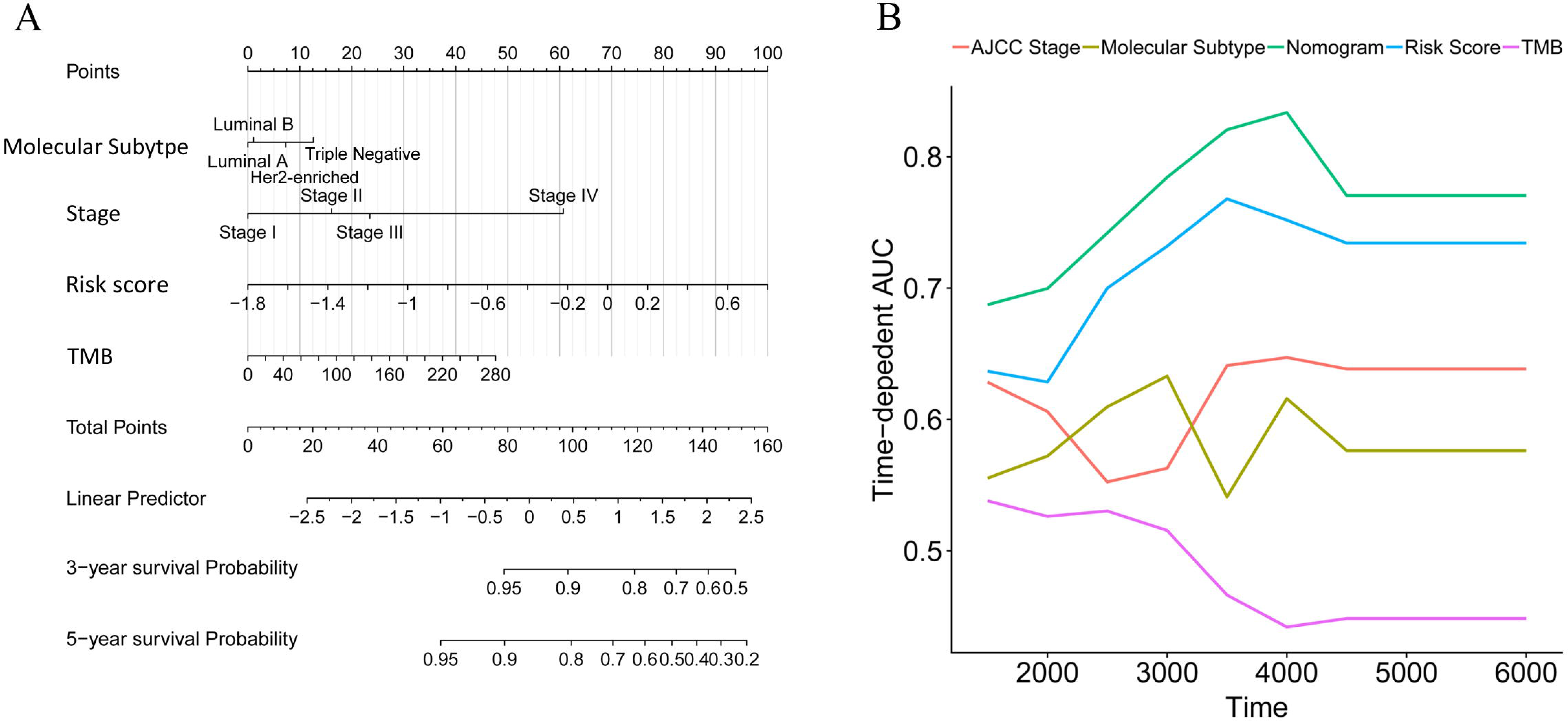
Construction of a nomogram for survival prediction. A, Nomogram combining the immune signature with clinicopathological features. B, The AUC(t) of the multivariable models indicated that the nomogram had the highest predictive power for overall survival.

### The immune signature predicted the immunotherapeutic benefits in IDC patients

VEGF-A, the main mediator in tumour angiogenesis, hinders T cell infiltration in the tumour microenvironment. Hence, we explored the correlation between VEGF-A expression and the T cell immune response in IDC tumours. Interestingly, the increased VEGFA expression significantly correlated with both decreased levels of activated CD8+ T cells and Th1 cell infiltration in the high immune infiltration tumour microenvironment but not in the low immune infiltration tumour microenvironment (Fig. 8A and 8B). Furthermore, perforin, the molecular effector found in the granules of cytotoxic T lymphocytes and natural killer cells, also showed a negative correlation with VEGF-A expression (Fig. 8C). Finally, the positive correlation of VEGF-A and the risk score was identified. PD-1, PDL-1 and cytotoxic T lymphocyte antigen-4 (CTLA-4) are promising targets for the treatment of patients with breast and non-small cell lung cancer. PD-1, PDL-1, and CTLA-4 antibodies are undergoing studies for the treatment of breast cancer. We analysed the correlation of PD-1, PDL-1, and CTLA-4 expression in the high- and low-infiltration groups. The expression of PD-1, PDL-1, and CTLA-4 was more significantly correlated with CD8+ T cells, Th1 cell ssGSEA score and perforin expression in the high-infiltration group than in the low-infiltration group. Furthermore, the mean expression of PD-1, PDL-1, and CTLA-4 was significantly increased in the high-infiltration group, indicating a potentially enhanced response to the corresponding anticancer antibody for IDCs with high immune infiltration status. In our constructed immune signature, the risk score showed a negative correlation with PD-1, PDL-1, and CTLA-4 expression, which implies a potentially enhanced effect of PD-1, PDL-1, and CTLA-4 antibodies in patients with low risk score. Lastly, we checked the correlation of the expression profiles of several immune checkpoint proteins, e.g., CD160, CD274, CD276, CTLA-4, LAG3, and PDCD1, risk score, and VEGF-A in the TCGA and GSE20685 cohorts (Fig. S8).

**Fig 8.**
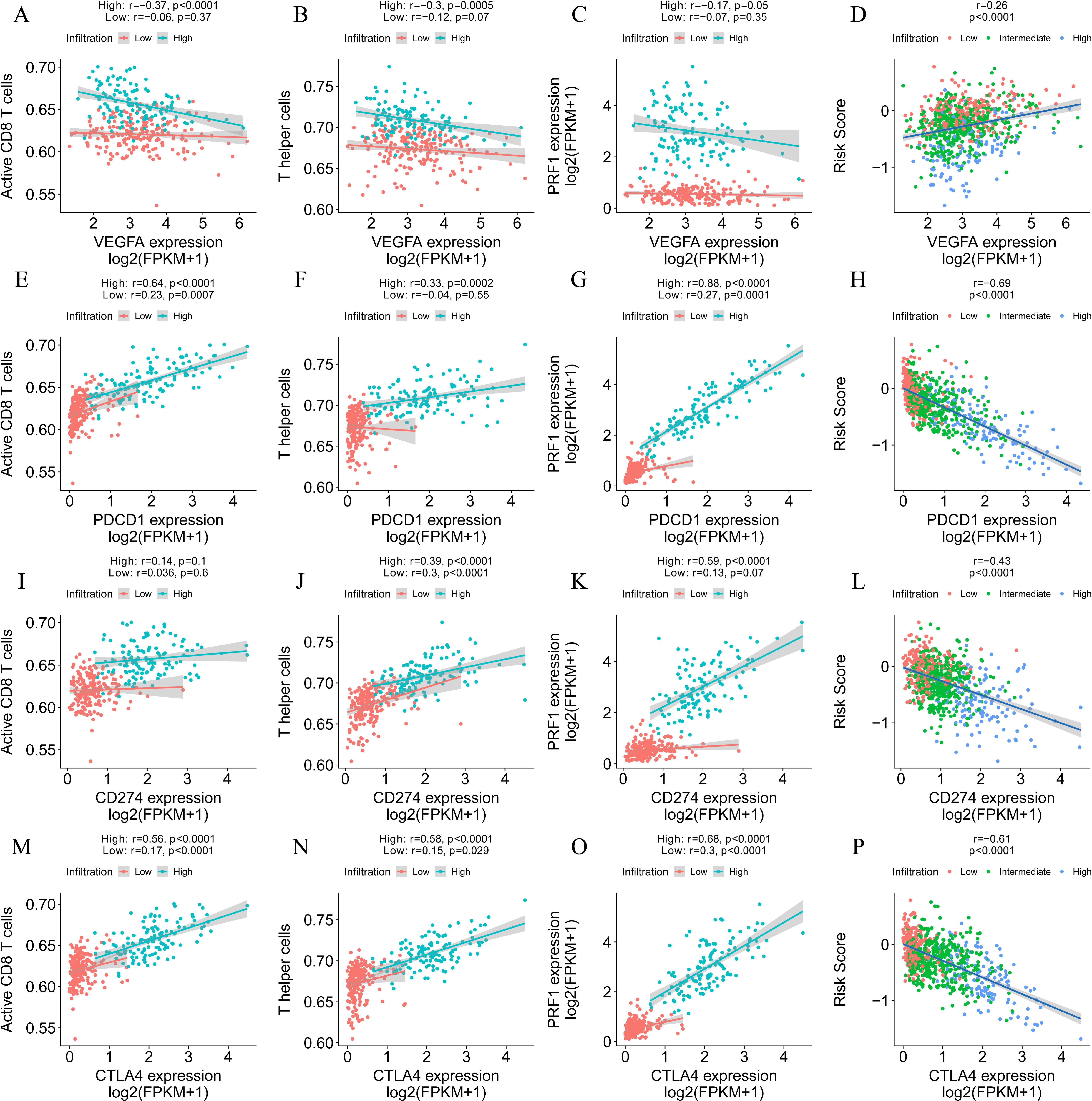
Immune signature predicts immunotherapeutic benefits. A, B and C, The correlation of VEGFA expression with T cell infiltration, Th1 cells and PRF1 expression in high and low immune infiltration conditions. D, The correlation of VEGFA expression with the immune signature. E, F and G, The correlation of PD-1 expression with T cell infiltration, Th1 cells and PRF1 expression in high and low immune infiltration conditions. H, The correlation of PD-1 expression with the immune signature. I, J and K, The correlation of PDL-1 expression with T cell infiltration, Th1 cells and PRF1 expression in high and low immune infiltration conditions. L, The correlation of PDL-1 expression with the immune signature. M, N and O, The correlation of CTLA-4 expression with T cell infiltration, Th1 cells and PRF1 expression in high and low immune infiltration conditions. P, The correlation of CTLA-4 expression with the immune signature.

## Discussion

In this study, we depicted the immune landscape of IDC using a large cohort. The immune landscape might explain the differences in prognoses of patients with IDC and responses to PD1, PDL-1 and CTLA-4 antibodies. Based on the immune landscape, we constructed an immune signature that calculated the risk score per patient. The correlation of signature and immune landscape revealed that the T cell-mediated immune response played a crucial role in the signature. Patients with low risk scores had increased T cell infiltration scores, interferon-γ signatures, and cytotoxic activity scores, indicating active T cell immune responses and favourable survival probability. A random forest algorithm was applied to find the most important somatic mutation correlated with the immune signature. A nomogram was constructed based on the immune signature and other clinicopathological properties of IDCs. A time-dependent ROC analysis showed high accuracy of the immune signature and nomogram in terms of predicting the survival of IDC patients. Lastly, PD-1, PDL-1, and CTLA-4 expression was found to be highly associated with the risk score. The patients with low risk scores had increased expression levels of PD-1, PDL-1, and CTLA-4, indicating a potentially high response rate to PD-1, PDL-1, and CTLA-4 antibodies.

In our analysis, the IDCs were clustered into three main clusters (low immune infiltration, intermediate immune infiltration, and high immune infiltration). The patients in the high-infiltration cluster had the best survival probability compared with patients in the low- and intermediate-infiltration clusters. The T cell immune response is the central event in antitumour immunity (14). T cells are divided into CD4+ (helper T cells, Th) and CD8+ (cytotoxic T cells, Tc) T cells. Their secretomes include IFN-γ, TNF-α, and IL17, which have antitumour effects. Hence, the increased T cell infiltration score, interferon-γ signature, and cytotoxic activity score may lead to an anti-tumour effect in the high-infiltration group. This finding could explain the different OS and RFS in the high- and low-infiltration groups.

From the immune landscape in IDCs, we built an immune signature that included seven features (QRSL1, TIMM8A, IGHA1, BATF, KLRB1, SPIB, and FLT3LG). FLT3LG is a crucial cytokine that controls the development of DCs and is particularly important for CD8-positive classical DCs and their CD103-positive tissue counterparts. A clinical trial is currently underway to treat melanoma patients with a combination of immunostimulatory FLT3LG and a peptide-based vaccine targeting DCs (15). KLRB1, which encodes CD161, a surface marker on several T cell subsets and NK cells, has been found to be most frequently associated with favourable outcomes in many cancer types by enhancing innate immune characteristics (16). SPIB is a member of the ETS family and profoundly affects B cell functions. B cells that lack SPIB fail to proliferate in response to IgM cross-linking, exhibit limited capacity to respond to T-dependent antigens, and produce low levels of IgG1, IgG2a, and IgG2b (17). In addition, SPIB can activate enhancer elements in both Ig-λ and Ig-κ genes, increasing the expression of these two genes. BATF is an inhibitor of AP-1-driven transcription. Recent studies have revealed that BATF can regulate positive transcriptional activity in dendritic cells, B cells and T cells (18). BATF leucine zipper motifs interact with interferon-regulatory factor 4 (IRF4) and IRF8 at AP-1–IRF consensus elements (AICEs), adding additional flexibility to the actions of IRF4 and IRF8, which were previously considered to interact with SPIB and PU.1 (19). The interaction of IRF4 and BATF in T helper 17 cells increases the production of IL-17, IL-21, IL-22 and IL-23 receptor. TIMM8A is involved in the import and insertion of hydrophobic membrane proteins from the cytoplasm to the mitochondrial inner membrane. The Bax/Bak complex mediates the release of DDP/TIMM8a and activates Drp1-mediated fission to promote mitochondrial fragmentation and subsequent elimination during programmed cell death (20). From the expression profiles of the seven genes above, we calculated the risk score for each patient and predicted the survival of IDC patients.

The risk score from the immune signature was most significantly correlated with the ssGSEA score of cytotoxic cells, CD8 T cells and T cells, indicating the important roles of the T cell immune response in the immune signature. Interestingly, DCs in the low-risk group played a more important role than DCs in the high-risk group. The increased proportion of DCs significantly correlated with favourable survival in the low-risk group but did not correlate with favourable survival of patients in the high-risk group. Th innate inflammatory cytokines, such as IL-1, IL-12, and IL-23 expressed by DCs, promote IFN-γ-secreting CD4+ T cell and cytotoxic T lymphocyte responses (21). The high proportion of DCs and T cells cooperate to achieve the antitumour effect in IDC patients with low risk scores. Furthermore, the GSEA results revealed high levels of IFN-γ, TNF-α, and TNF-α secretion in the low-risk group, which contribute to the antitumour activity in IDC patients with low risk scores. WGCNA revealed opposing directions of the risk score (cor = −0.64) and immune infiltration (cor = 0.8) with the brown module, indicating the high level of correlation of risk score (calculated by immune signature) and immune infiltration. The hub gene in the brown module plays an essential role in regulating immune infiltration. The GO analysis revealed that T cell activation was the most significantly enriched biological process, indicating that the T cell-mediated immune response is the central event in both immune infiltration and the immune signature.

The spectrum of somatic mutations varied in IDC patients. The different mutation burdens in IDCs led us to analyse whether the landscape of immune cells and the immune signature were associated with somatic mutations. The TMB showed a positive correlation with the risk score in IDC patients. Furthermore, a random forest algorithm was performed to identify the most important variables correlated with the immune signature. TP53, SCN10A, PIK3CA and 32 other genes were the most significant variables in the analysis. TP53 and PIK3CA mutations are the most common gene mutations in IDCs (44% and 33%, respectively). In the 35 gene variables, GATA3, a key regulator of ER activity, is a newly identified gene that is mutated in IDCs (5% in ILC versus 13% in IDC, q = 0.03) (3). Mutations in GATA3 are more frequent in luminal A IDC and are mutually exclusive with FOXA1 events. The differential expression level and enrichment for mutations of GATA3 in IDCs and of FOXA1 in ILC indicates a preferential requirement for the distinct regulation of ER activity in ILC and IDC (3). Previous studies revealed that the GATA3 mutation correlates with increased expression, which is associated with the immune response (22, 23). Our analysis further confirms the correlation of the GATA3 mutation with immune infiltration. In addition, we constructed a nomogram that integrated clinicopathological features with the immune signature to predict the survival probability of IDC patients. Compared with other clinicopathological features, the immune signature showed the best accuracy in predicting the survival of IDC patients at any time point and would therefore be helpful for the diagnosis and precise treatment of IDC patients.

There have been several studies for the treatment of breast cancer with immunotherapeutic antibodies. PD-1 is expressed by exhausted T cells. PD-1 and PD-L1 exhibit inhibitory receptor–ligand interactions, which are involved in the negative regulation of T cell activation and peripheral tolerance during immune responses by cancer cells. Despite demonstrated successes, only a proportion of patients benefit from PD-1 and PDL-1 antibody treatment. Hence, it is important to determine the mechanism that leads to the varied therapeutic effect of PD-1 and PDL-1 antibody treatment and thus improve individual diagnosis and precision medicine. PD-L1 expression, microsatellite instability and deficient mismatch repair are important biomarkers that predict the response to anti-PD-1/PD-L1 therapies (24–26). Among the three biomarkers, PD-L1 expression has been validated in nearly all tumour types for all approved anti-PD-1/PD-L1 therapies. In our analysis, the expression of PD1, PDL-1, and CTLA-4 was significantly increased in the high-infiltration group. Furthermore, the expression of PD1, PDL-1, and CTLA-4 had a significant correlation with CD8+ T cells, Th1 cell ssGSEA score and perforin expression in the high-infiltration group, which provides a basis for PD-1/PD-L1 and CTLA-4 treatment. Similarly, the immune signature we constructed also indicated that high expression levels of PD1, PDL-1, and CTLA-4 correlated with low risk score. Therefore, patients with a low risk score could derive more benefit from immunotherapy than patients with a high risk score.

In the current study, we performed a comprehensive evaluation of the immune landscape of IDC and constructed an immune signature related to the immune landscape. This analysis of TME immune infiltration patterns has shed light on how IDC respond to immune checkpoint therapy and may guide the development of novel drug combination strategies.

## List of abbreviations

TME: tumour microenvironment
IDC: invasive ductal carcinoma
TCGA: The Cancer Genome Atlas
GEO: Gene Expression Omnibus
ssGSEA: single-sample gene set enrichment
LASSO: least absolute shrinkage and selection operator
ILC: invasive lobular carcinoma
DEG: differentially expressed gene
WGCNA: weighted correlation network analysis
TMB: total mutation burden
IRF4: interferon-regulatory factor 4
AICE: AP-1–IRF consensus element.

## Declarations

### Ethics approval and consent to participate

Not applicable

### Consent for publication

Not applicable

### Availability of data and materials

The datasets supporting the conclusions of this article are available in the Xena browser (https://xenabrowser.net/) repository.

### Author Contributions

XW. B. and R. S. conceived and designed the experiments. XW. B., S. X., and K. Z. analysed the data. XW. B., and YF. W. wrote the paper. YB. Z., K. Z., and R. S. reviewed the draft. All authors read and approved the final manuscript.

## Acknowledgements

We would like to thank Dr Michael Rosemann for helpful discussions and suggestions.

## Figure legends

**Fig S1.**
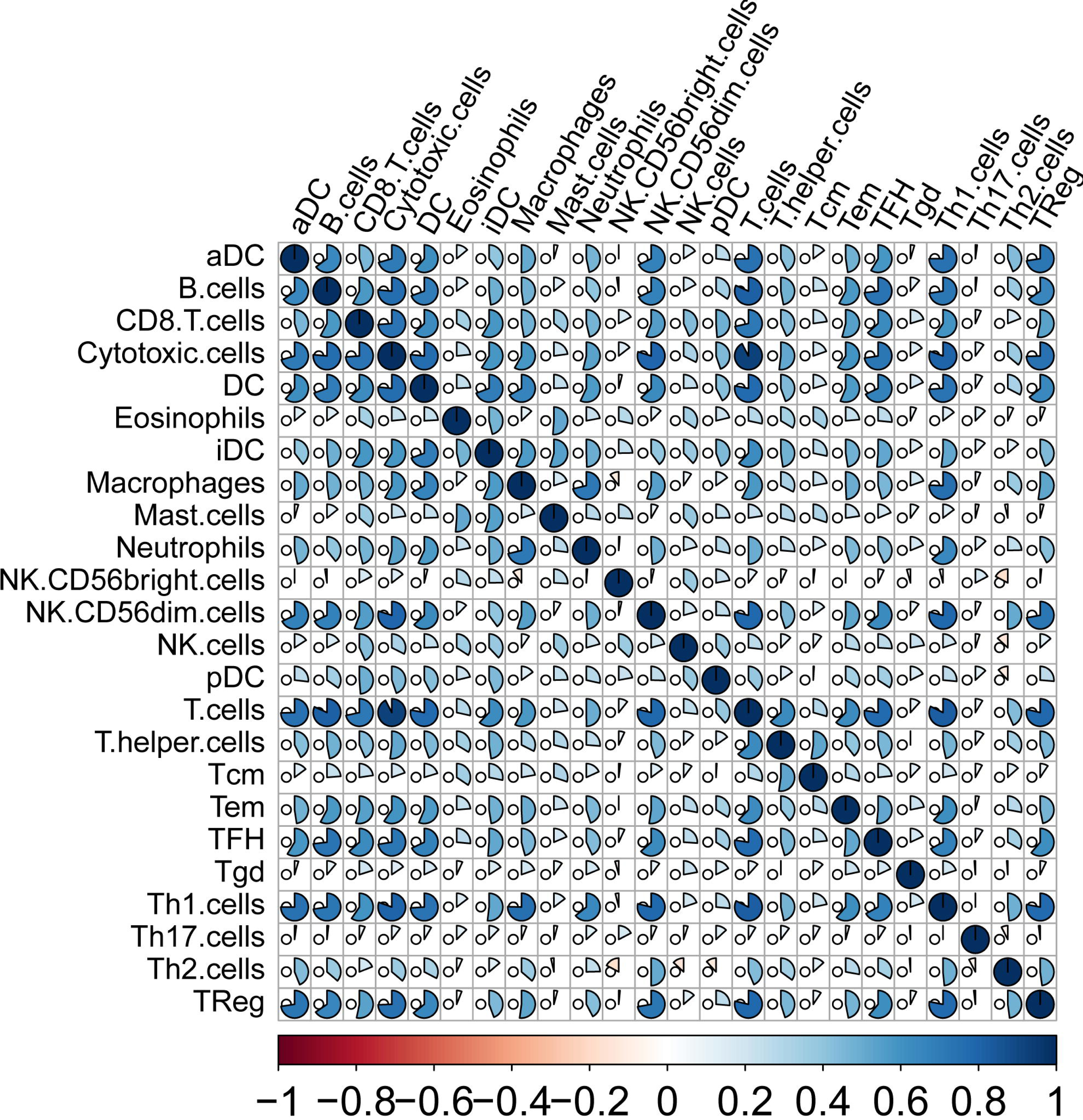
The correlation between different infiltrating immune cells.

**Fig S2.**
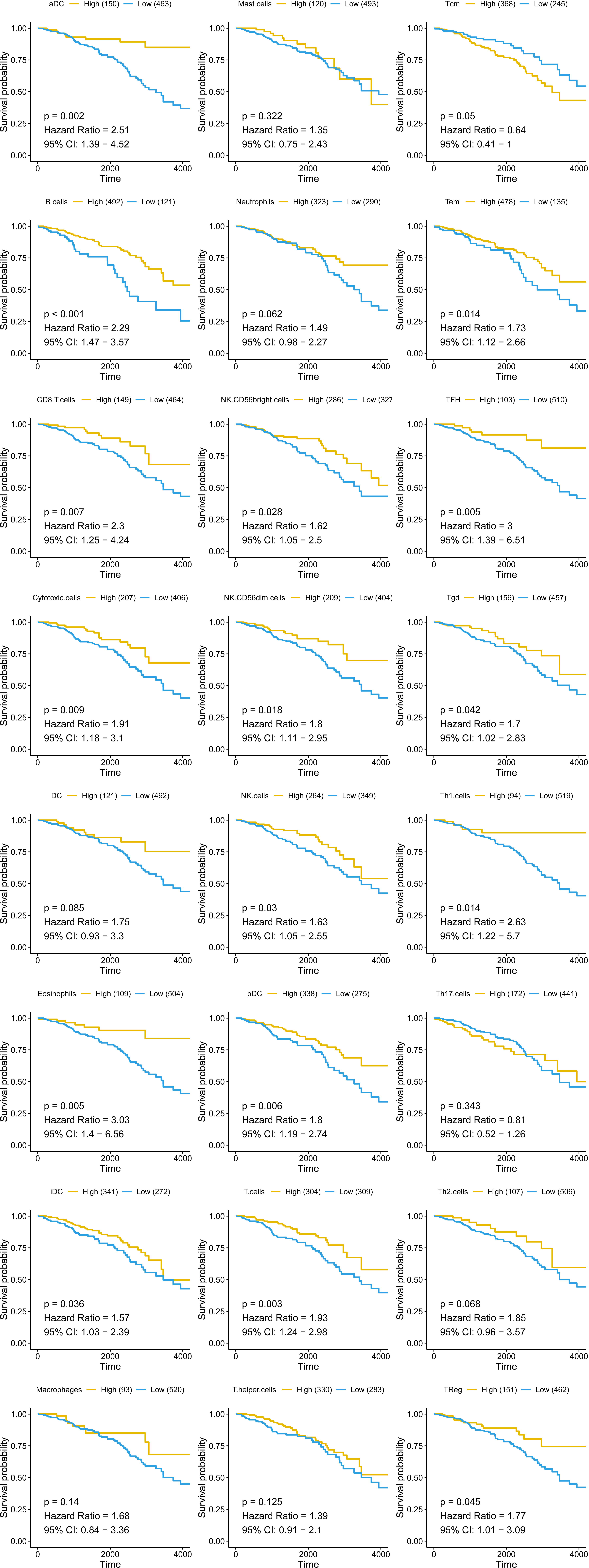
The correlation between the ssGSEA scores of infiltrating immune cells and the OS probability of IDC patients.

**Fig S3.**
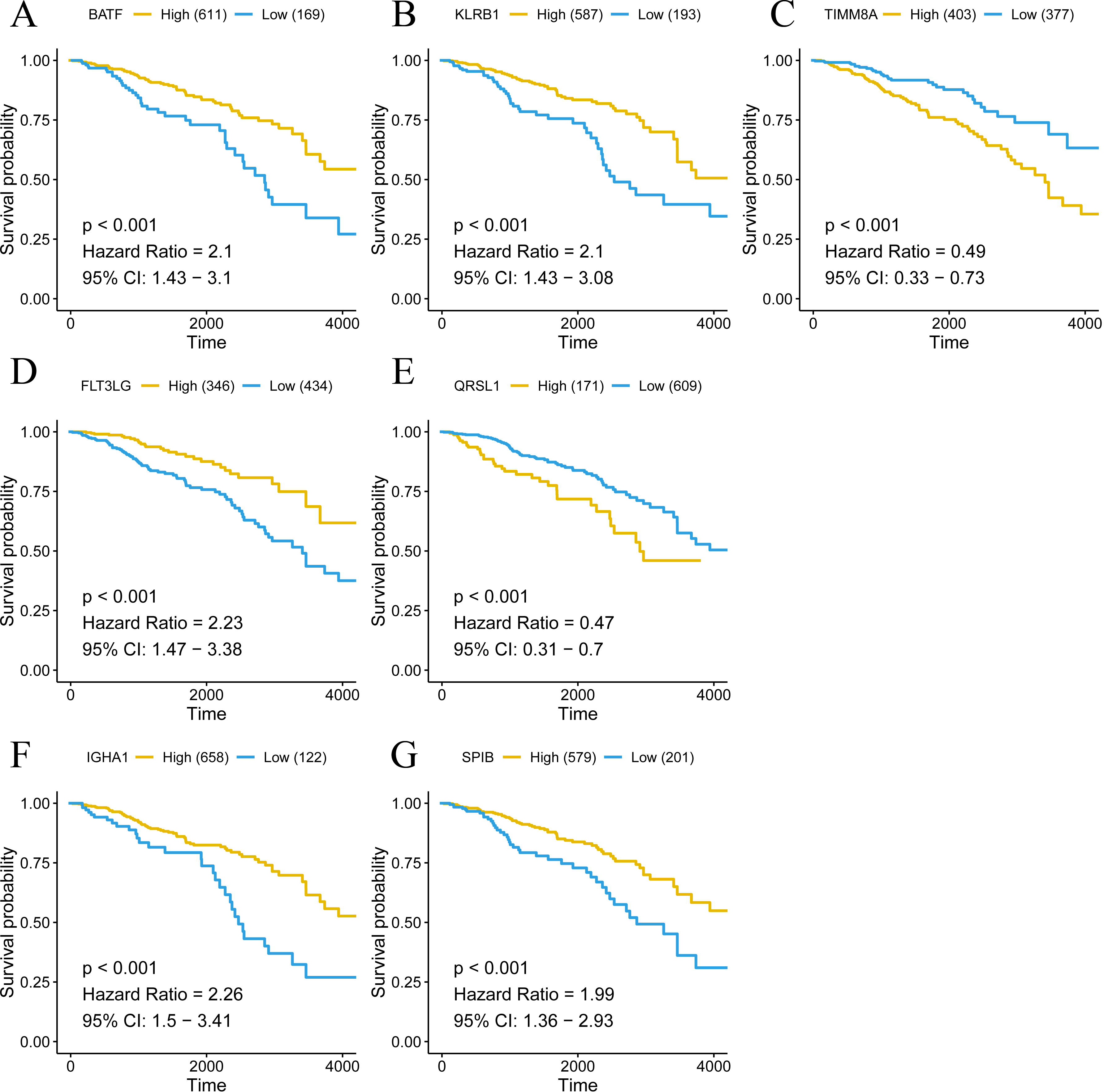
The correlation between the expression level of seven genes in the immune signature and the OS probability of IDC patients.

**Fig S4.**
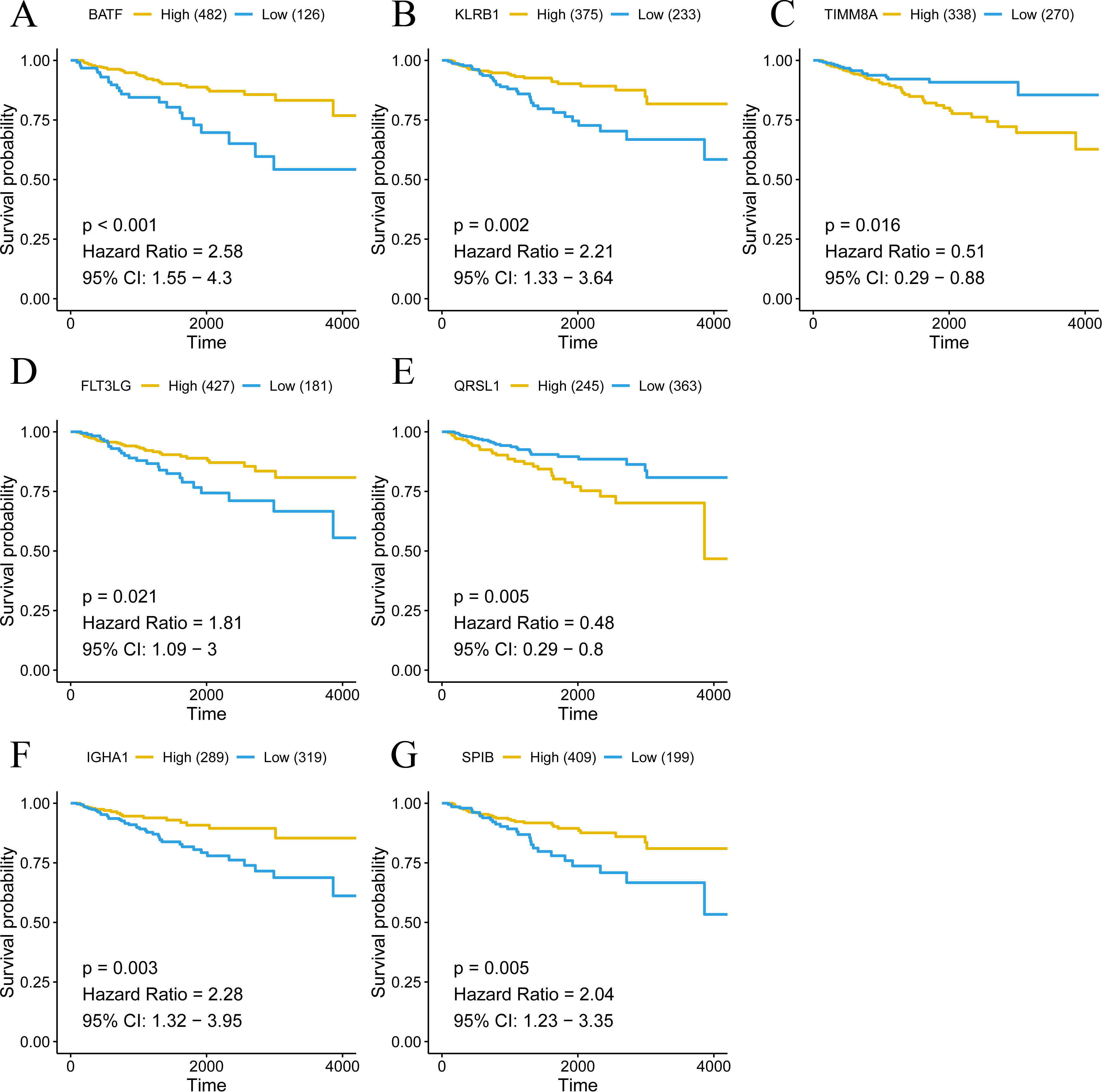
The correlation between the expression of seven genes in the immune signature and the RFS probability of IDC patients.

**Fig S5.**
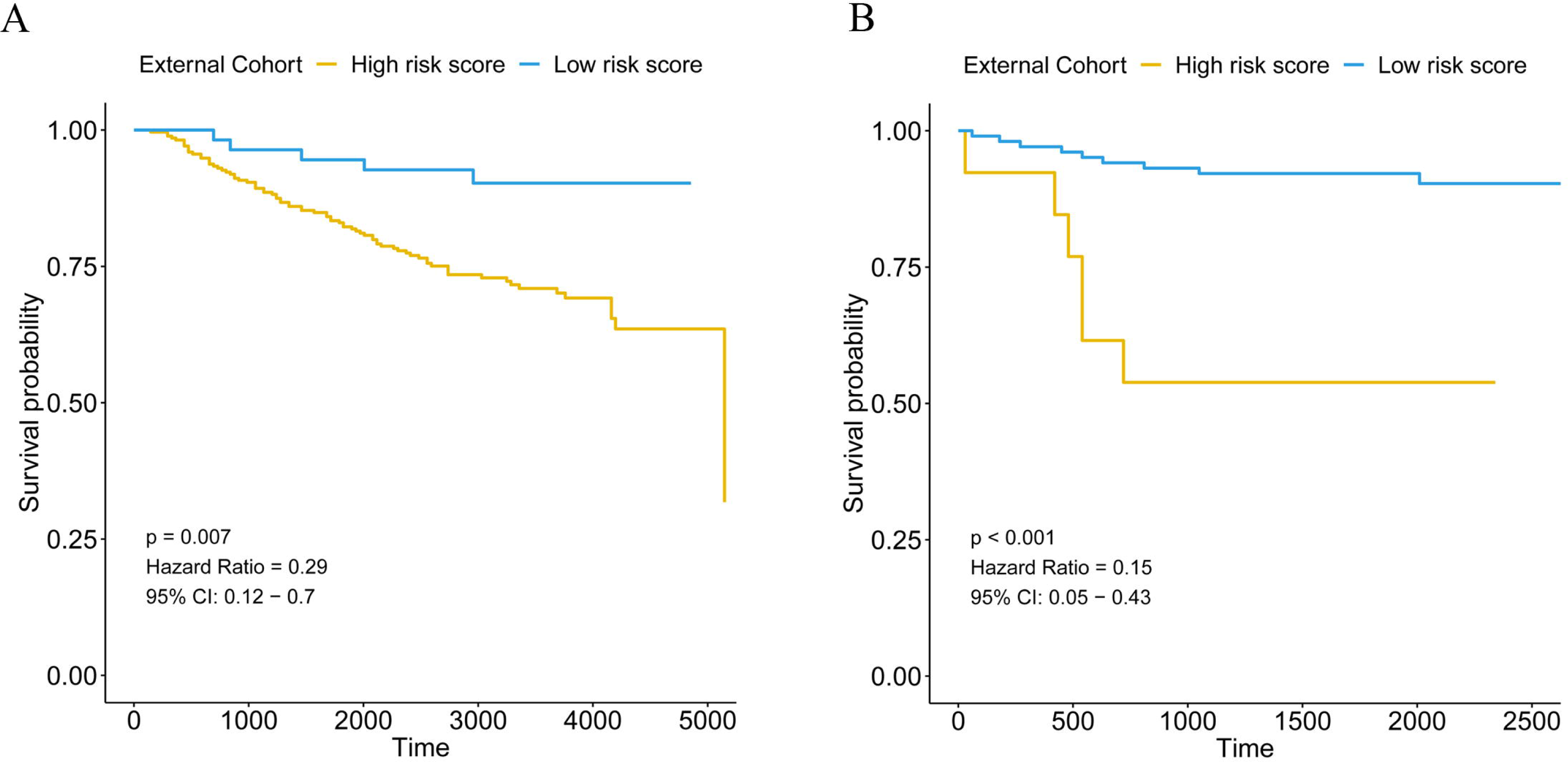
Validation of the immune signature in two external cohorts, GSE20685 (A) and GSE86948 (B).

**Fig S6.**
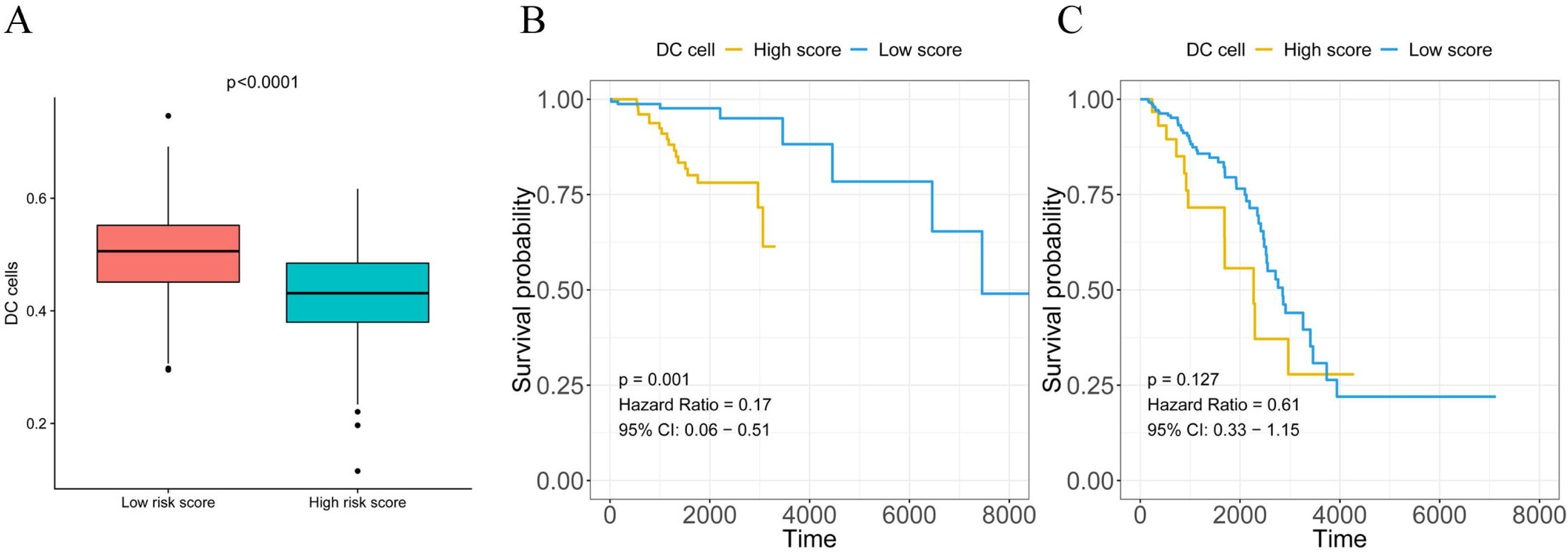
The correlation between the ssGSEA scores of DCs and the OS probability of IDC patients in the high- and low-risk score groups. A, The ssGSEA scores were higher in the high- and low-risk score groups. B, The correlation between the ssGSEA scores of DCs and the OS probability of IDC patients in the low-risk score group. C, The correlation between the ssGSEA scores of DCs and the OS probability of IDC patients in the high-risk score group.

**Fig S7.**
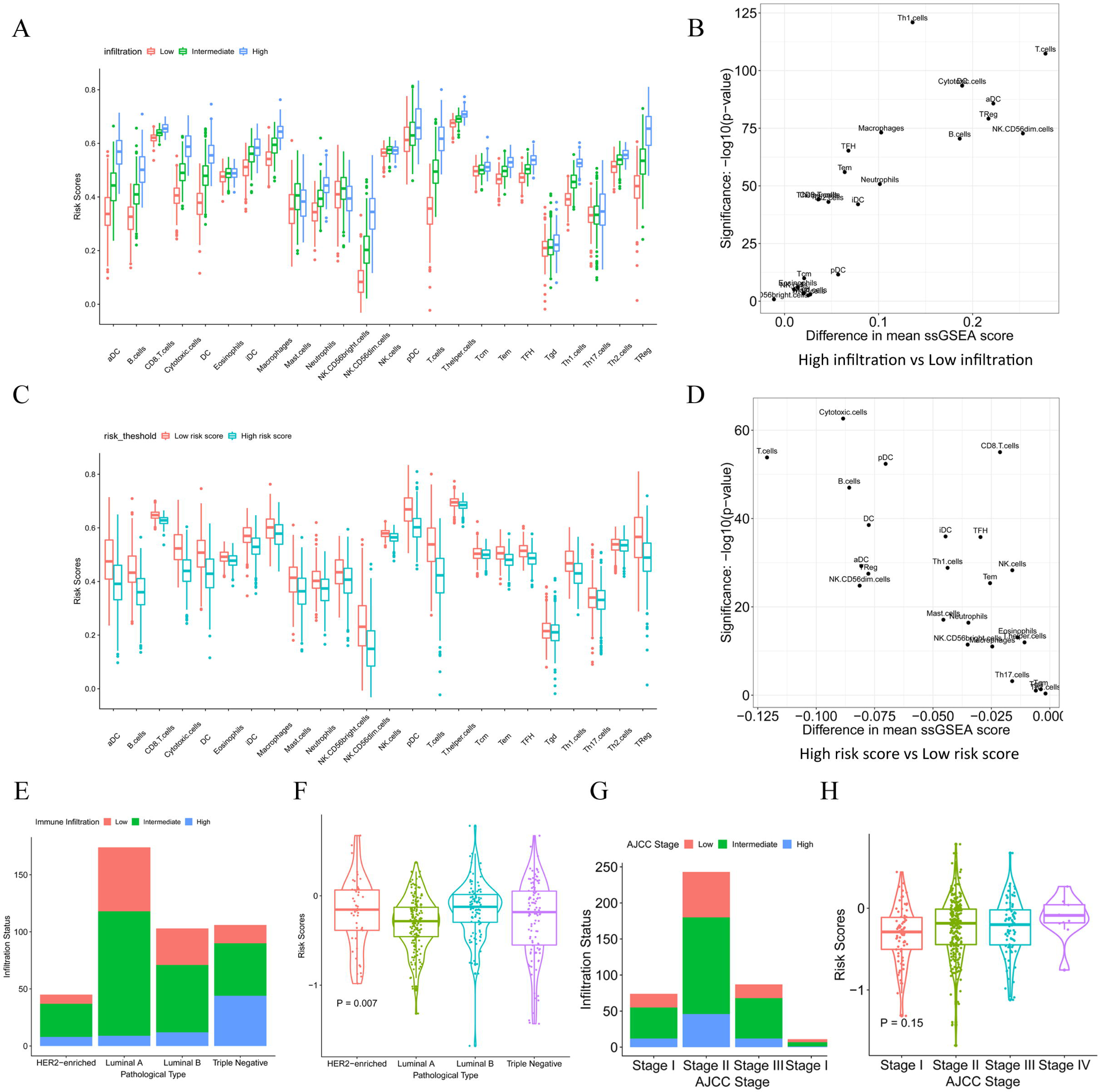
The ssGSEA score distribution in the low, intermediate and high immune infiltration patterns and in the low- and high-risk score groups. A, The ssGSEA score distribution in low, intermediate and high immune infiltration patterns. B, The difference and P value from the comparison between the ssGSEA score from low and high immune infiltration patterns. C, The ssGSEA score distribution in the low- and high-risk score groups. D, The difference and P value from the comparison between the ssGSEA score from the low- and high-risk score group. E, The distribution of immune infiltration patterns in different pathological subtypes. F, The distribution of risk scores in different pathological subtypes. G, The distribution of immune infiltration patterns at different pathological stages. H, The distribution of risk scores at different pathological stages.

**Fig S8.**
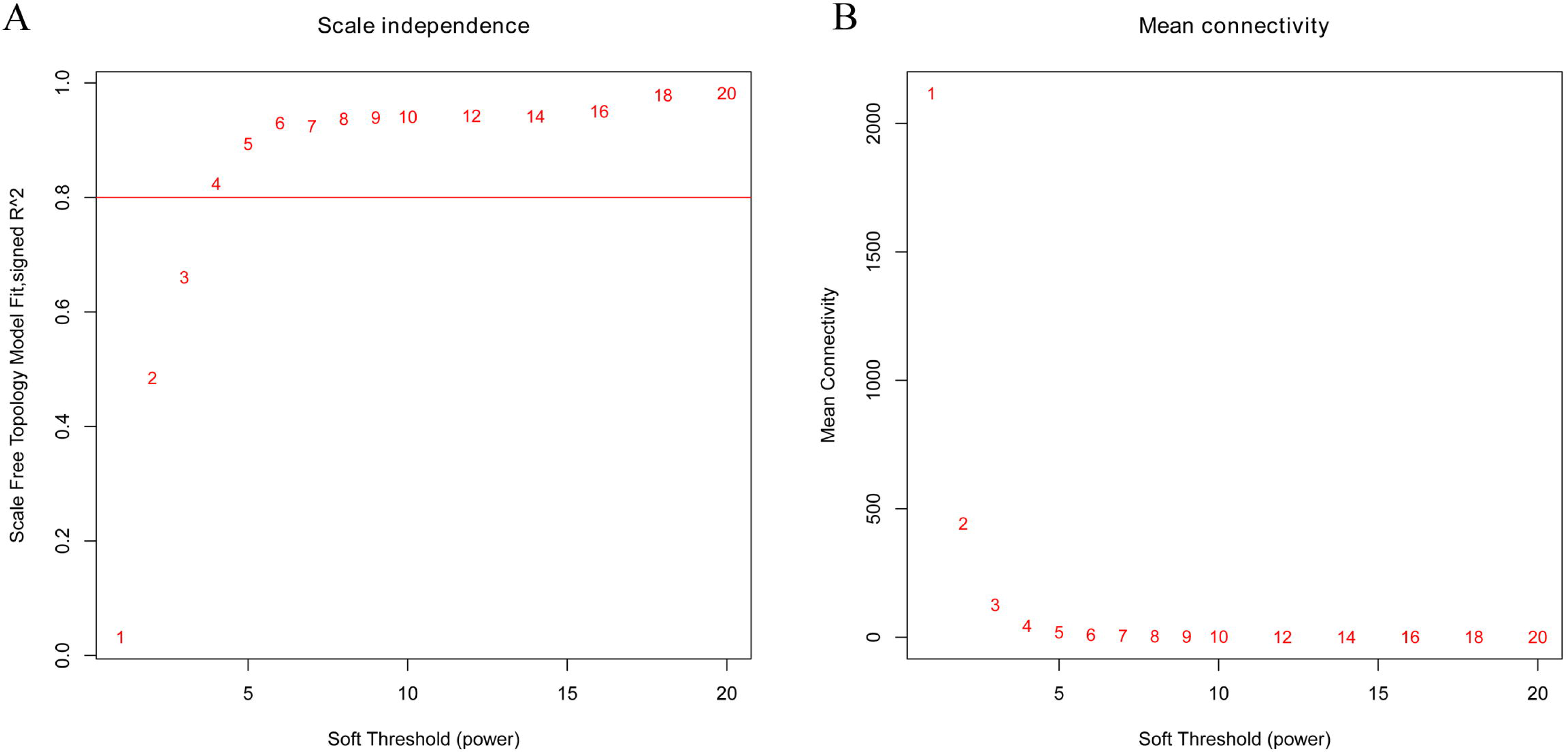
The selection of the soft threshold in the WGCNA.

**Fig S9.**
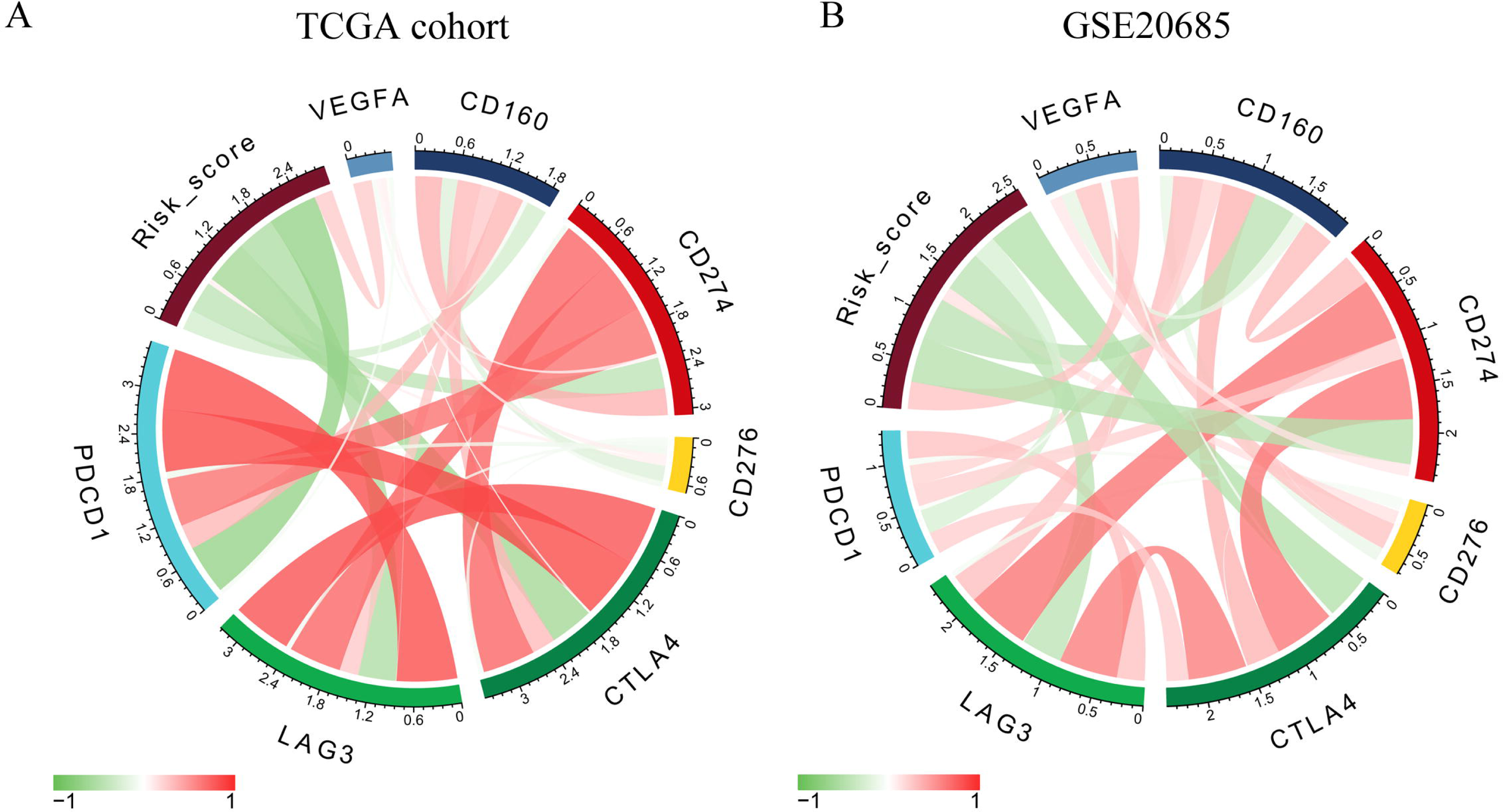
The correlation of the expression profiles of several immune checkpoint proteins, risk score, and VEGF-A in the TCGA (A) cohort and GSE20685 cohort (B).

